# Insight into the mechanism of molecular interactions of the NDM metallo-beta-lactamase and cyclic boronate inhibition by molecular docking and molecular dynamics simulations

**DOI:** 10.64898/2026.09.09.750529

**Authors:** Cui Shi, Yi-Feng Shi

**Affiliations:** School of Biological Engineering, Dalian Polytechnic University, Dalian 116034, China

**Keywords:** Antibiotic resistance, Metallo-β-lactamases, Carbapenems, Cyclic boronate inhibitor, Molecular docking, Molecular dynamics simulation, Molecular interactions

## Abstract

The spread and evolution of antimicrobial resistance (AMR) pose significant threats to public health, food safety, and sustainable development. Metallo-β-lactamases (MBLs), a class of carbapenemases that hydrolyze carbapenems—often considered last-resort antibiotics—are of particular concern and underscore the urgent need to develop effective inhibitors. Recent advancements in bicyclic boronate inhibitors represent significant progress toward this goal. Although the mechanistic basis of NDM-1 inhibition by bicyclic boronate inhibitors remains incompletely elucidated, this study systematically characterizes the binding profiles and intermolecular interaction landscapes between NDM-1 and four cyclic boronic acid-based inhibitors—vaborbactam (RPX7009), taniborbactam (VNRX 5133), xeruborbactam (QPX7728), and ledaborbactam (VNRX-5236). We established the NDM-1-mediated hydrolysis of two representative carbapenem antibiotics, meropenem and imipenem, as the reference system for comparative analysis. While conventional docking workflows often struggle to model metalloenzyme active sites accurately, the AMDock (Assisted Molecular Docking) platform, integrated with AutoDock’s optimized zinc force field, enables robust and reliable docking simulations for this enzyme family. Our AMDock-based molecular docking not only identified the critical NDM-1 binding-site residues governing hydrogen-bond networks with ligands, but also revealed that bicyclic boronate inhibitors show substantially greater binding stability than both carbapenems and their monocyclic boronate counterparts. A follow-up 100 ns all-atom molecular dynamics simulation in GROMACS provided atomistic-level evidence for this enhanced performance: RMSD, RMSF, radius of gyration, continuous hydrogen-bonding occupancy, and 2D free-energy landscape analyses all consistently supported the higher binding affinity of bicyclic boronates. Ultimately, this study characterizes two promising drug candidates, QPX7728 and VNRX-5133, as competitive inhibitors that can effectively rescue carbapenem activity against NDM-1 MBL-mediated antimicrobial resistance. This visualization elucidates interactions between NDM-1 and its two ligand classes—carbapenem antibiotics and cyclic boronate inhibitors. It reveals the stability and interaction strength underlying the distinct inhibition pattern of bicyclic boronate inhibitors in the enzyme’s active site, offering a promising strategy to combat antibiotic resistance.

## Introduction

Antimicrobial resistance (AMR) is a growing global health crisis driven by misuse and overuse of antibiotics in humans, animals, and plants, leading to the emergence of resistant pathogens that pose challenges in treating infections. ^[1]^ β-lactam antibiotics account for more than half of injectable antibiotics worldwide. Bacteria can develop resistance to β-lactams in several ways, including producing β-lactamase enzymes. β-Lactamases are broadly categorized as serine β-lactamases (SBLs) and metallo-β-lactamases (MBLs) based on their catalytic mechanism. These enzymes efficiently catalyze the hydrolytic opening of the β-lactam ring, preventing the antibiotic from targeting PBPs. One response to this threat is combining β-lactam antibiotics with β-lactamase inhibitors, which are used in the clinic to overcome resistance by inhibiting β-lactamases.

β-lactam antibiotics include penicillins, cephalosporins, carbapenems, penems (also known as thiopenems), and monobactams. This classification depends on the chemical nature of the ring fused to the β-lactam pharmacophore unit, generating a non-coplanar bicyclic scaffold. Emerging bacterial resistance has limited their antibacterial efficacy. Carbapenems are the latest generation of β-lactam antibiotics and, as such, are currently used as last-resort drugs in Intensive Care Units. Thus, among β-lactamases, carbapenemases represent the main scourge in the clinics ^[2–3]^.

Beta-lactamases are divided into four classes based on primary sequence homology and differences in hydrolytic mechanisms: A, B, C, and D ^[4]^. Classes A, C, and D β-lactamases are serine enzymes, which can hydrolyze the β-lactam ring via a serine-bound acyl intermediate in the active site, whereas class B β-lactamases (named metallo-β-lactamases, MBLs) present one or two zinc ions in the active site, which are necessary for their enzymatic activity ^[5–6]^. However, the continued use of β-lactams, which is often excessive and inappropriate, has led to the spread of resistance to β-lactams, to extended-spectrum cephalosporins (e.g., cefotaxime, ceftriaxone, and ceftazidime), and more recently to carbapenems (doripenem, ertapenem, imipenem, meropenem). MBLs are unique in their unusually broad substrate spectrum, being able to hydrolyze penicillins, cephalosporins, carbapenems, and even β-lactam-based SBL inhibitors such as clavulanic acid and sulbactam, i.e., all classes of bicyclic β-lactams. This makes MBLs unique because they can efficiently hydrolyze carbapenems, thus representing a challenge for the clinical use of these antibiotics. As a result, the resistance conferred by these enzymes cannot be countered. However, a varied set of promising compounds that abolish their activity have been identified, some of which are currently in clinical trials ^[7–8]^.

The global effort to prevent β-lactam resistance focuses on developing broad-spectrum β-lactamase inhibitors that mimic the β-lactam core and block β-lactamases, including cephalosporinases and serine-based carbapenemases, that severely limit antimicrobial activity. Boronic acids are rapidly emerging as highly promising scaffolds for cross-class β-lactamase inhibition. Bicyclic boronates (BCBs) such as taniborbactam (TAN) and others are capable of simultaneously inhibiting both SBLs and MBLs. This is desirable because no MBL inhibitors approved for clinical use, and bacterial strains co-expressing SBLs and MBLs are increasingly encountered ^[9–10]^.

Integrating computational approaches such as molecular docking, quantitative structure-activity relationship (QSAR) modeling, and molecular dynamics (MD) simulations can significantly improve the identification and optimization of potential natural product inhibitors against NDM-1^[11–12]^. Molecular docking simulations help explore binding interactions between these compounds and the NDM-1 enzyme, providing insights into binding affinities and modes of action. By evaluating the stability and dynamics of protein-ligand complexes over time, molecular dynamics simulations further refine these findings and help identify inhibitors that are both effective and robust. Previous studies have used molecular docking ^[13]^ and molecular dynamics simulations to explore the binding mechanism between intact β-lactam compounds and New Delhi metallo-β-lactamase 1 (NDM-1) ^[14–15]^. Given that the mechanistic basis underpinning NDM-1 inhibition by bicyclic boronate inhibitors remains incompletely resolved, the present study systematically characterizes the binding profiles and intermolecular interaction landscapes of NDM-1 with three representative bicyclic boronate agents—taniborbactam (VNRX-5133), xeruborbactam (QPX7728), and ledaborbactam (VNRX-5236). This analysis uses head-to-head comparative profiling with two clinically deployed carbapenem antibiotics, meropenem and imipenem, as well as the licensed monocyclic boronate β-lactamase inhibitor vaborbactam (RPX7009). While molecular docking techniques have reached a considerable level of maturity, the resulting protein-ligand complex models typically only account for protein flexibility within a localized scope. In this context, molecular dynamics simulations are critical for validating docking conformations and investigating intermolecular interactions in protein-ligand complexes with full intrinsic flexibility.

## 1. Materials and methods

### 1.1 Collection and Preparation of ligands and proteins

The three-dimensional structures of NDM-1 (PDB ID: 4EYL), NDM-1 bound to hydrolyzed meropenem (PDB ID:5YPN) ^[16]^, NDM-1 bound to hydrolyzed imipenem (PDB ID:5YPI) ^[16]^, NDM-1 bound to VNRX-5133 (PDB ID:6RMF)^[17]^ and NDM-1 bound to QPX7728 (PDB ID:6V1M)^[18]^ were obtained from the RCSB Protein Data Bank (https://www.rcsb.org/). The native structures of the ligands were retrieved from the PubChem database (https://pubchem.ncbi.nlm.nih.gov/)^[19]^ in SDF format (.sdf) which was subsequently converted into the PDB format in PyMOL (Version 2.5).

Before molecular docking, we prepared the target protein in PyMOL. This involved removing the second chain, assigning bond orders using default settings, and removing all attached ligands and water molecules (except the catalytic ones). We also carefully preserved the integrity of the zinc ions and the catalytic water within the protein structure.

### 1.2 Molecular Docking

The molecular docking workflow was implemented on AMDock^[20]^ using the AutoDock4Zn algorithm, with the AMBER force field applied and the system pH set to 7.4 to mimic physiological conditions. Meropenem (PubChem CID: 441130), as a reference drug, is co-crystallized with metallo-beta-lactamase (PDB: 4EYL), allowing us to define the binding site of the enzyme. We preprocessed the ligand in OpenBabel for full geometry optimization and atomic charge calculation before docking. We centered the docking box on the original ligand position in the crystal structure to cover the entire binding pocket, keeping all other parameters at the software’s default settings. After docking completion, the optimal conformation with the highest predicted binding stability was selected and saved in PDB format for further analysis.

### 1.3 Molecular dynamics simulations

The molecular docking complex with the top-ranked binding energy among all protein-ligand systems was selected for subsequent molecular dynamics simulations. The docked ligand conformation was first visually inspected in PyMOL and exported as a properly formatted mol2 file. The GROMACS corresponding GAFF-topology for the small molecule was then generated using the Sobtop program and RESP charges derived from quantum calculations using Multiwfn can be directly incorporated into the ligand topology^[21]^.

We used GROMACS 2022.4 to perform molecular dynamics simulations of protein-ligand complexes ^[22]^. During the simulation, we used the Amber99sb force field and the TIP3P water model for solvation. The protein-ligand complex was placed in a dodecahedral water box with a minimum 1.0 nm distance between the protein surface and box boundary, and Na⁺/Cl⁻ counterions were added to neutralize the system’s net charge. The system underwent a two-stage energy minimization protocol: 5000 steps of steepest descent relaxation with a force ceiling of 1000 kJ·mol⁻¹·nm⁻¹, followed by conjugate gradient minimization until all atomic residual forces fell below 1000 J·mol⁻¹·nm⁻¹ for full convergence. Before productive sampling, we subjected the initial configuration to two consecutive equilibration runs. We first equilibrated the system in the NVT ensemble for 1000 ps (2 fs time step), ramping the temperature from 0 K to 300 K, then performed a 1000 ps NPT equilibration at 300 K and 1 atm to stabilize temperature and pressure and achieve equilibrium density. Finally, a 100 ns molecular dynamics simulation was performed at constant temperature (300 K) and pressure (1 atm) with a 2 fs time step.

### 1.4 Complex structure stability characterization and visual mapping

The backbone stability and conformational dynamics of the simulated protein were analyzed by calculating the Root Mean Square Deviation (RMSD), Root Mean Square Fluctuation (RMSF) and Radius of Gyration (Rg) using the built-in analysis tools of GROMACS, with all results visualized for subsequent structural assessment. We used gmx hbond to quantify ligand-protein hydrogen-bond dynamics, exported results as .xvg files, and visualized them in QtGrace. Protein backbone RMSD and radius of gyration (Rg) from MD trajectories were preprocessed. We calculated the 2D free energy landscape, projected onto RMSD (x-axis) and Rg (y-axis), using the native gmx sham module in GROMACS. 2D ligand–protein interaction plots were generated with LigPlot, and 3D binding pose visualizations were rendered in PyMOL.

## 2. Results

### 2.1 Molecular Docking

**Fig 1.**
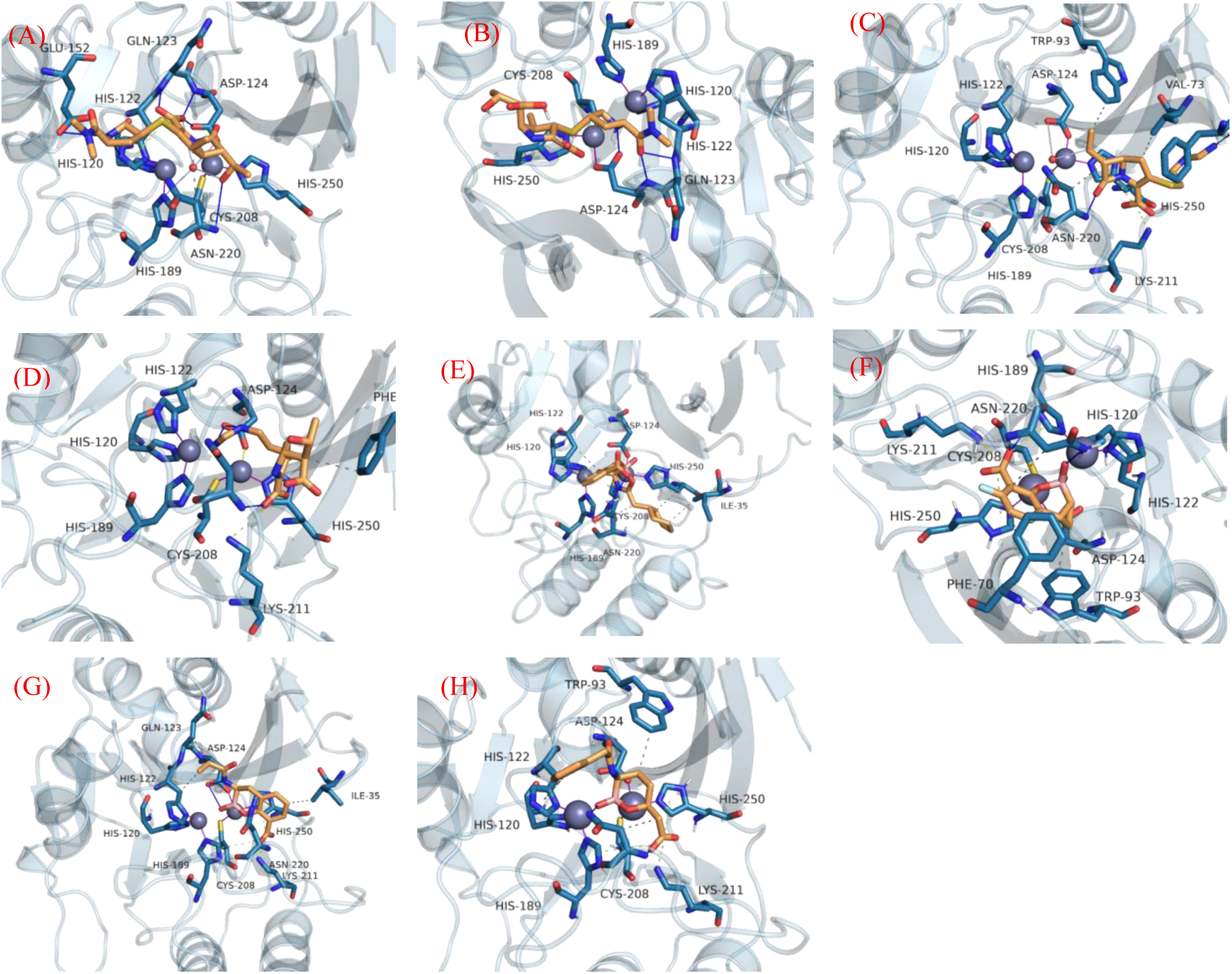
3D Diagram of molecular interaction between NDM-1 and its ligands (A) Meropenem;(B) Hydrolyzed meropenem;( C) Imipenem; (D) Hydrolyzed imipenem (E) VNRX-5133; (F) QPX7728; (G) VNRX-5236; (H) RPX7009

We performed molecular docking simulations in AMDock to characterize the binding profiles of four cyclic boron-based inhibitors (RPX7009, QPX7728, VNRX-5133, VNRX-5236) against NDM-1, with pre- and post-hydrolysis imipenem and meropenem as reference controls. All ligands were docked into the NDM-1 active pocket, and the corresponding binding poses and affinity data are presented in Fig. 1 and Table 1.

**Table 1.**
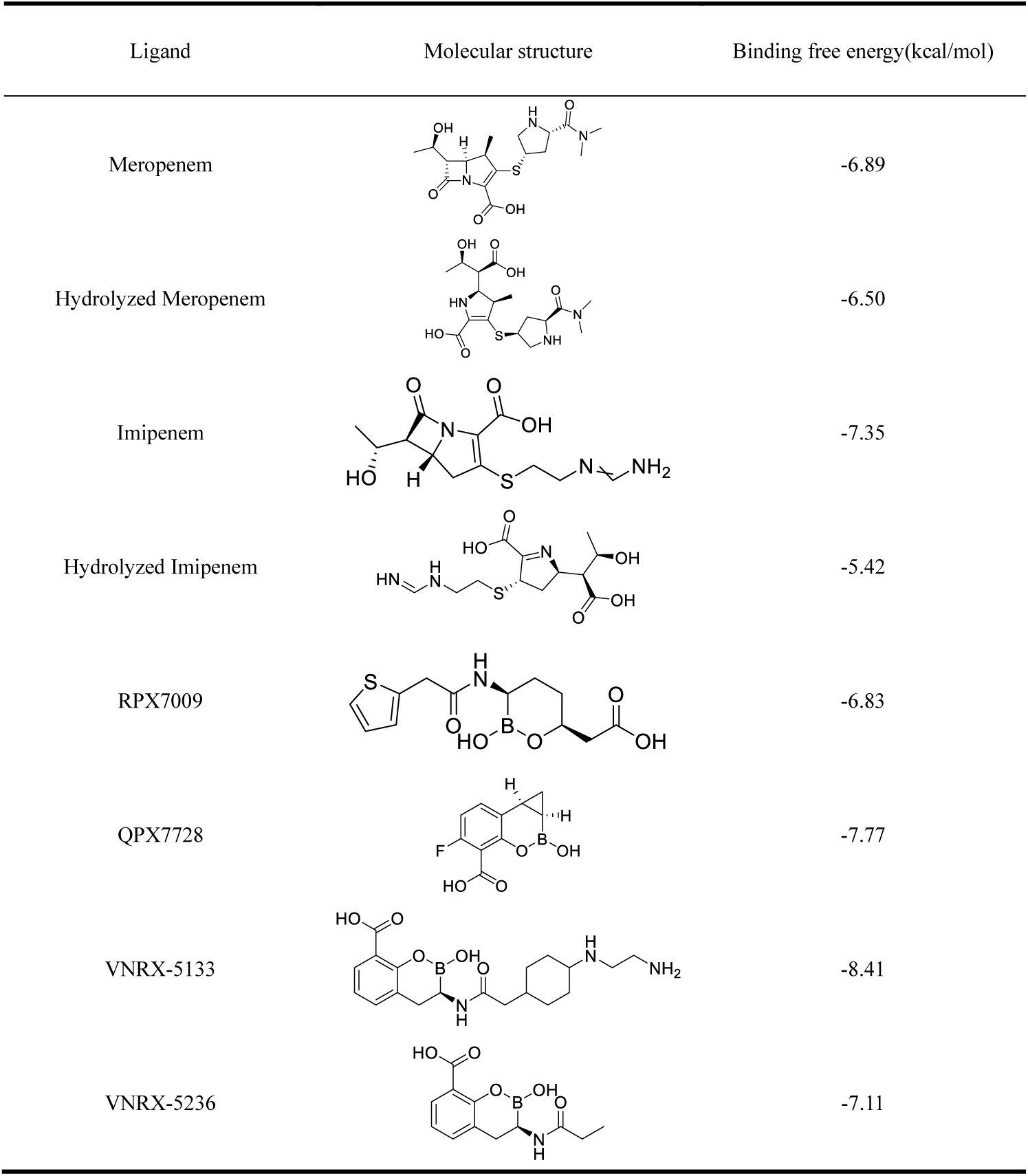
Binding free energy of NDM-1 with different ligands in Molecular Docking.

| Ligand | Molecular structure | Binding free energy(kcal/mol) |
| --- | --- | --- |
| Meropenem |  | -6.89 |
| Hydrolyzed Meropenem |  | -6.50 |
| Imipenem |  | -7.35 |
| Hydrolyzed Imipenem |  | -5.42 |
| RPX7009 |  | -6.83 |
| QPX7728 |  | -7.77 |
| VNRX-5133 |  | -8.41 |
| VNRX-5236 |  | -7.11 |

For the molecular docking simulation of NDM-1, a typical di-zinc metalloprotein, the conventional docking workflow often fails to accurately maintain the coordination environment of metal ions and the key water-mediated interaction network at the active site. To address this limitation, this study adopted the AMDock program, which integrates dedicated, pre-calibrated zinc-ion force field parameters. During structure preprocessing, we did not remove the conserved water molecule between the two catalytic Zn²⁺ ions in the NDM-1 crystal structure, as in standard procedures. Instead, we explicitly retained this catalytically critical water molecule and converted it into the standard PDBQT format with the protein structure, ensuring it was not excluded in the subsequent docking calculation. This optimized protocol finally generated a reliable NDM-1-ligand binding mode, in which the active-site water molecule directly participates in the hydrogen bond and coordination interaction between the protein and the small-molecule ligand, effectively improving the simulation fidelity compared with the conventional water-free docking strategy.

To date, no crystal structure of NDM-1 bound to an intact antibiotic substrate has been resolved; only structures of hydrolyzed antibiotic-NDM-1 complexes are available. Here, we used molecular docking to characterize the pre-hydrolysis binding mode of NDM-1 with its substrates. AMD molecular docking results, as quantified by binding free energy from docking scoring, demonstrate that the hydrolytic products of meropenem and imipenem exhibit significantly weaker binding affinities toward NDM-1, with respective values of -6.50 kcal/mol and -5.42 kcal/mol, compared to their corresponding native substrates. Molecular docking was performed with a pre-fixed enzyme structure and conformation, without incorporating the induced-fit mechanism that allows dynamic conformational adjustment of the enzyme.

The binding free energies of all tested inhibitors are significantly more favorable than those of the two reference antibiotics. VNRX-5133 exhibits the highest binding affinity with a ΔG of -8.41 kcal/mol, followed by QPX7728 (-7.77 kcal/mol) and VNRX-5236 (-7.11 kcal/mol), while RPX7009 displays the lowest affinity at -6.83 kcal/mol. These computational results generally agree with the experimentally determined Ki values: RPX7009 (Ki > 40 mM), VNRX-5133 (Ki = 0.081 mM), and QPX7728 (Ki = 0.032 mM). This finding demonstrates that the bicyclic boronic acid inhibitors QPX7728, VNRX-5133, and VNRX-5236 possess superior inhibitory properties compared to the monocyclic boronic acid inhibitor RPX7009^[17–18]^.

**Fig 2.**
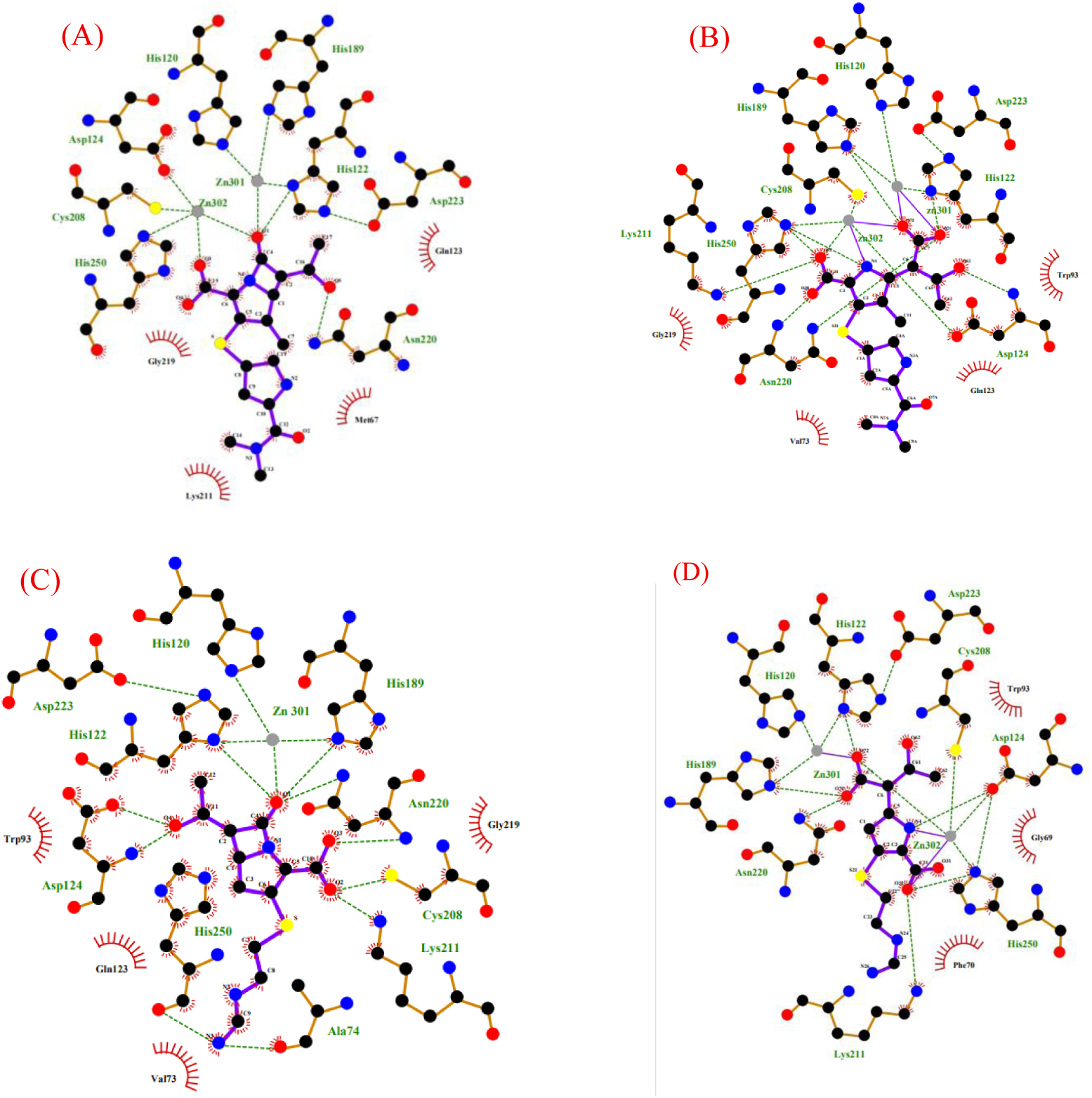
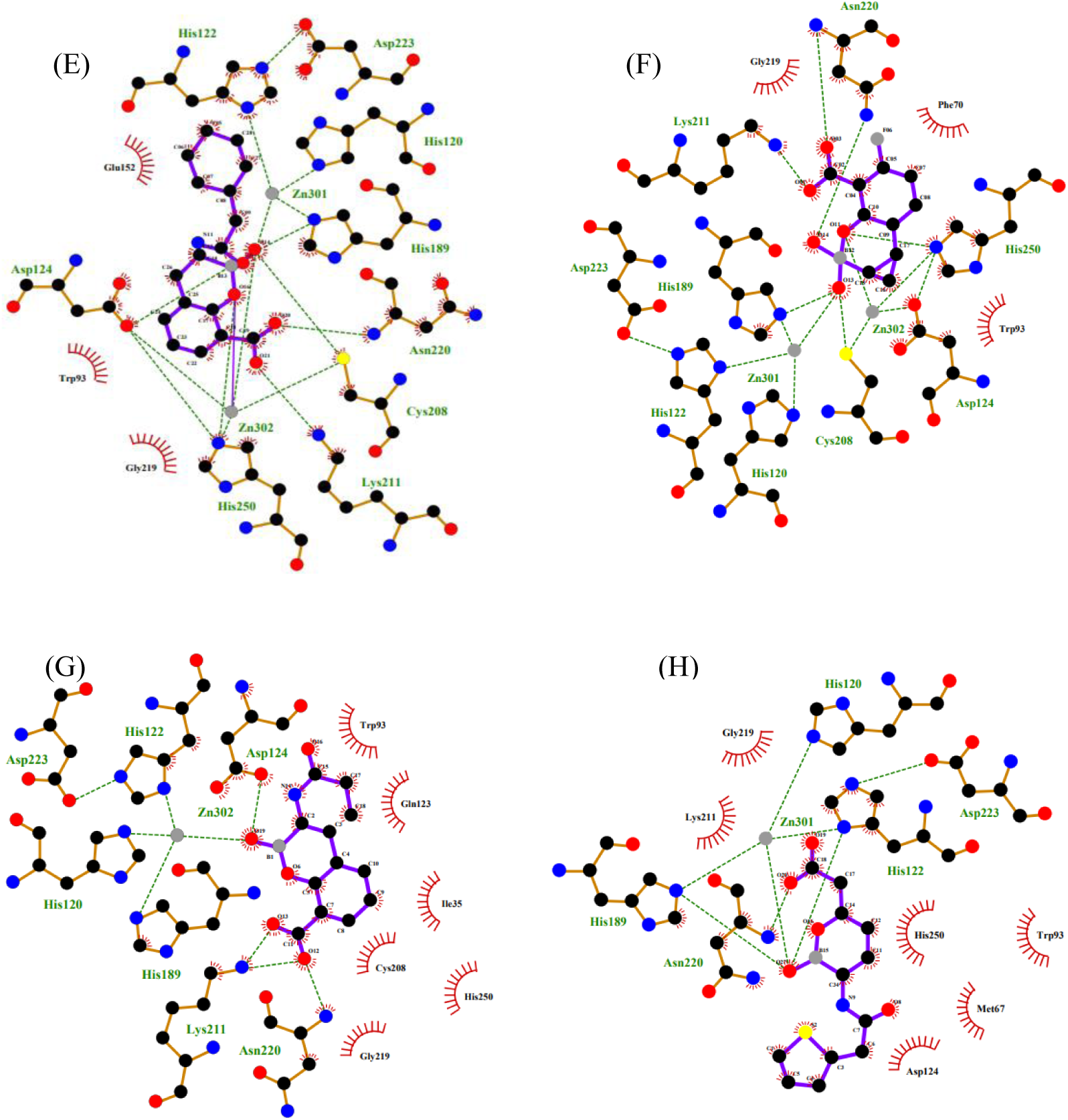
2D diagram of molecular interactions between NDM-1 and its ligands (A) Meropenem;(B) Hydrolyzed meropenem;( C) Imipenem; (D) Hydrolyzed imipenem (E) VNRX-5133; (F) QPX7728; (G) VNRX-5236; (H) RPX7009 (The green dashed line represents hydrogen bonds and metal coordination bonds with zinc ions; the red dashed line represents hydrophobic interactions)

Two-dimensional interaction mapping (Fig. 2) reveals that the inhibitor coordinates stably to the active-center zinc ion via its boronate group, alongside specific intermolecular interactions with proximal amino acid residues, which completely block NDM-1 activity.

### 2.2 Molecular Dynamics Simulation

**Fig 3.**
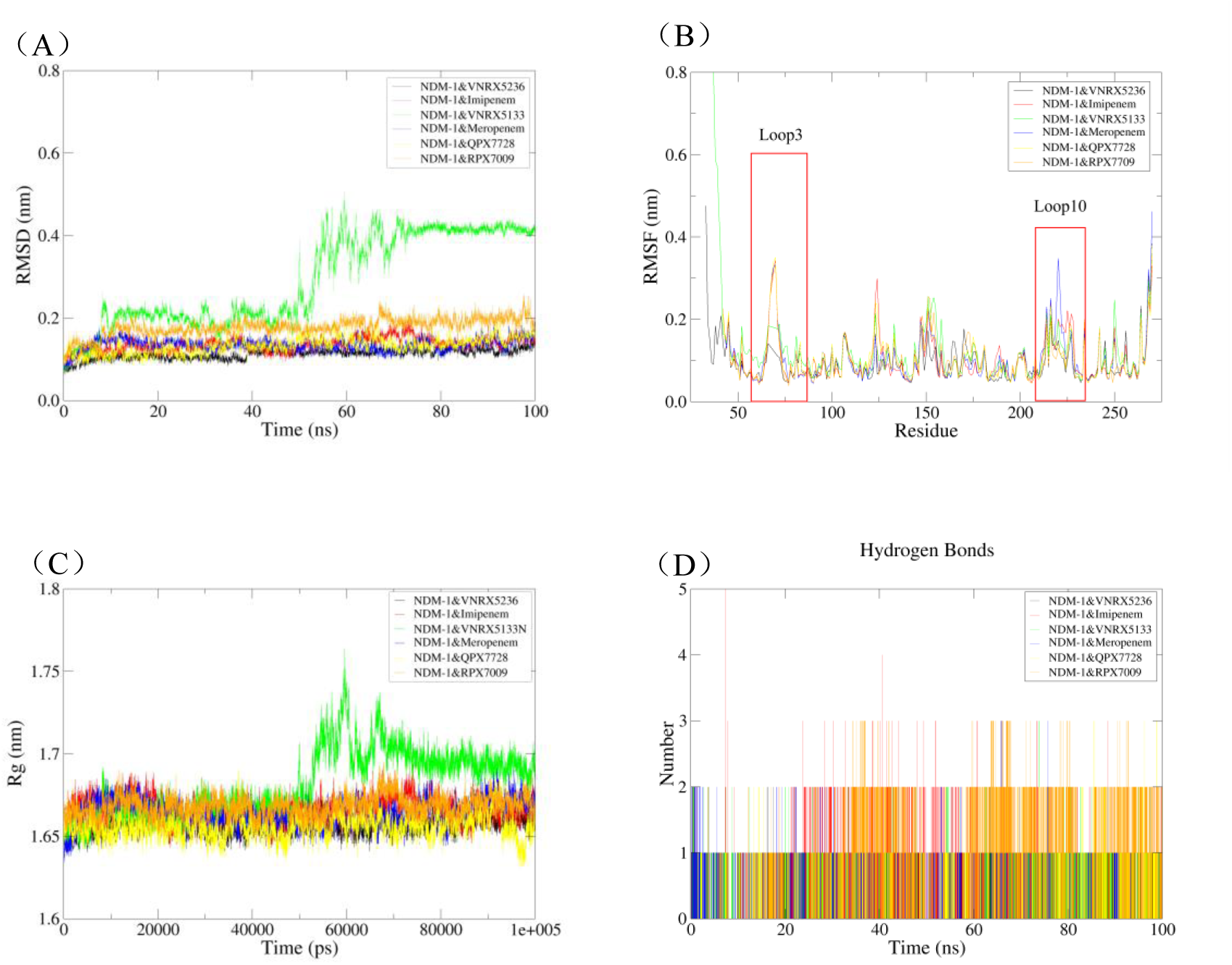
Molecular dynamics simulation results of the ligand-NDM-1 complex (A: RMSD; B: RMSF; C: Rg; D: hydrogen bonds)

To further evaluate the binding stability of the formed complex and the intermolecular interaction profiles, 100 ns molecular dynamics simulations were carried out on the docked ligand-protein complex via the GROMACS simulation package. A 100 ns molecular dynamics simulation trajectory captures key biological events, including local conformational rearrangements of the target protein, spontaneous ligand association and dissociation, and intrinsic loop relaxation dynamics, providing an optimal trade-off between simulation accuracy and computational overhead. We analyzed the simulated trajectories by calculating backbone RMSD, protein RMSF, radius of gyration (Rg), and hydrogen bonds formed between the ligand and target protein. Figure 3 presents the results.

In MD simulations, protein RMSD fluctuations indicate ligand-induced conformational changes or local structural instability, which may alter protein function and binding affinity. RMSD analysis is critical for studying ligand binding stability and mechanisms. As shown in Fig. 3A, all simulated systems, excluding the NDM-1-VNRX-5133 adduct, converged to a stable equilibrium state within 20–30 ns, with their RMSD values confined to a narrow fluctuation window of 0.08–0.16 nm. Compared with the NDM-1 antibiotic substrates, the inhibitors (VNRX-5236 and QPX7728) show lower RMSD values and more confined fluctuation amplitudes upon binding to NDM-1, indicating competitive inhibition. Notably, the bicyclic inhibitors VNRX-5236 and QPX7728, particularly VNRX-5236, show lower RMSD and less conformational fluctuation than both the substrate and the monocyclic inhibitor RPX7009.

RMSF analysis quantitatively describes the regulatory effects of ligands on the flexibility and rigidity of protein residues, which is critical for developing ligands, such as inhibitors, that can enhance protein stability. Figure 3B shows the 100 ns RMSF profile of NDM-1 amino acid residues. Apart from the high flexibility of unconstrained C- and N-termini, residues around 55–75, 125, 150, and 210–225 exhibit prominent fluctuations, corresponding to the flexible Loop3, Loop10, and α4 regions in NDM-1. Compared with the substrate-bound state, NDM-1 exhibits markedly reduced conformational flexibility in its amino acid residues upon inhibitor binding. This observation indicates that the inhibitor restricts flexibility in these loop regions, locks NDM-1 in an inactive conformation, and forms a tighter, more stable complex, explaining its induced-fit inhibition mechanism from a structural dynamics perspective.

Rg quantifies protein compactness and folding state, critical for studying folding, unfolding, oligomerization, and ligand-triggered global conformational changes. Figure 3C shows that all complexes except NDM-1/VNRX-5133 remain stably fluctuating at 1.64–1.69 nm, with high compactness and structural stability. NDM-1 shows large 60–75 ns fluctuations followed by stabilization when bound to VNRX-5133. This suggests the inhibitor likely induces conformational rearrangement in the flexible region adjacent to the active site to optimize binding, increasing the radius of gyration (Rg) of the whole protein.

The hydrogen bond count between protein and ligand directly determines ligand-receptor complex stability: more hydrogen bonds correspond to higher complex stability, making monitoring hydrogen-bond dynamics essential throughout molecular simulation. As shown in the MD simulation hydrogen-bond profile (Fig. 3D), the substrate imipenem shows a fluctuating hydrogen-bond count of 1–4, whereas the inhibitor maintains 1–3 hydrogen bonds with NDM-1, a lower value than that of the substrate. Despite the limited number of hydrogen bonds, their high occupancy ensures stable, long-lasting complex binding. Existing mechanistic studies show that a persistent hydrogen-bond network, when synergistically coupled with auxiliary non-covalent interactions such as hydrophobic contacts and π-π stacking, can markedly enhance the ligand’s conformational stability and target-binding affinity through multidimensional molecular recognition.

## 3. Discussion

3.1 The Structural Basis and Catalytic Mechanism of NDM-1 Metallo-β-lactamase X-ray spectroscopic data confirm that all MBLs possess an identical αβ/βα protein fold^[23–24]^. The Loop3 (Leu65–Val73), Loop7 (Thr119–Met126) and Loop10 (Cys208–Leu221) regions constitute the critical segments that shape the active site (Fig. 4). The active site of NDM-1 harbors two zinc ions. Zn1 is tightly coordinated by His120, His122, His189 and a water/hydroxyl moiety, adopting a tetrahedral geometry. Zn2 is coordinated by Asp124, Cys208 and His250, forming a trigonal pyramidal configuration. Zn1 and Zn2 form a dinuclear zinc cluster bridged by the Asp124 side chain. Catalytically, the two zinc ions position a hydrolytic water molecule for nucleophilic attack on the β-lactam carbonyl carbon, triggering cleavage of the carbapenem C7–N4 amide bond and consequent antibiotic deactivation (Fig. 5) [25–26].

**Fig 4.**
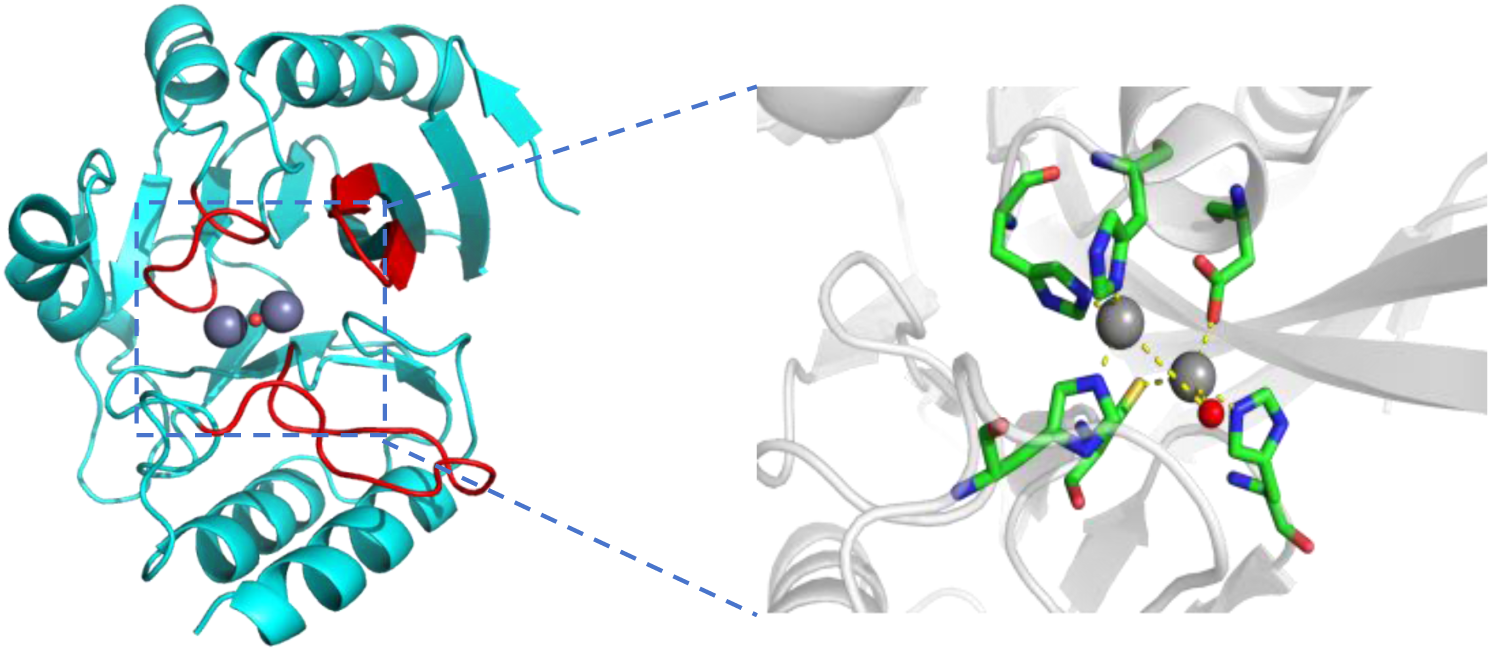
NDM-1 active site (gray spheres represent zinc ions, red spheres represent water molecules, and critical regions affecting the active site are marked in red)

**Fig 5.**
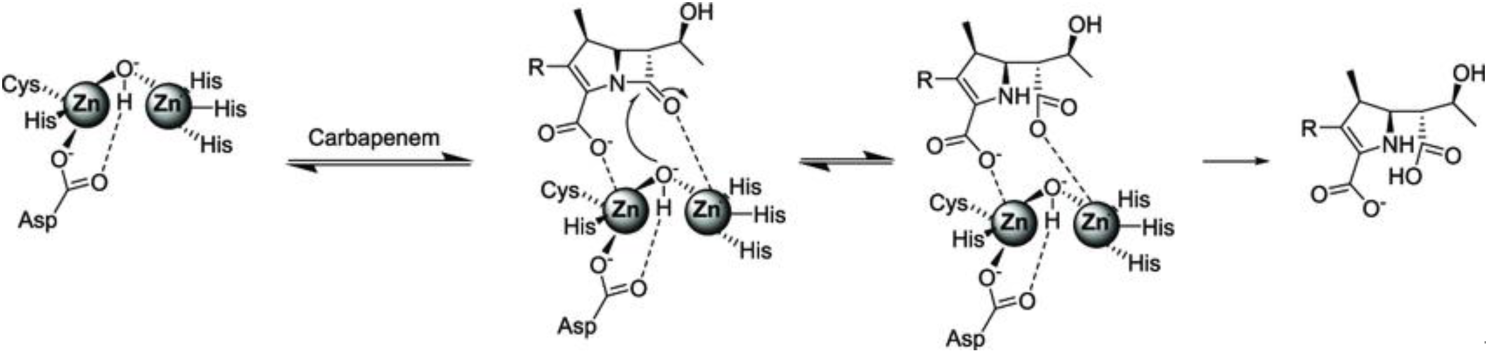
MBLs require the participation of active site metals (one or two Zn^2+^ ions in vivo) to activate an active site water molecule for ring opening hydrolysis of the β-lactam ring

### 3.2 Key Residues for NDM-1 Catalysis and Inhibitor Binding

Based on the “enzyme-structure-activity-inhibition” theoretical correlation, we further analyzed ligand-protein interactions. As shown in Table 2 and Fig. 6, the results reveal differences in the types and numbers of hydrogen-bond-forming amino acid residues in NDM-1-ligand interactions.

Asp124 and Asn220 form hydrogen bonds with nearly all ligands, marking them as key residues for active-site ligand recognition. As a direct zinc-coordinating residue, Asp124 is functionally more critical. In contrast to NDM-substrate antibiotics, their corresponding boronic acid-derived inhibitors establish hydrogen-bonding interactions with key residues including His189 and Cys208, as represented by the clinical-stage compounds QPX7728 and VNRX-5133. The participation of the side-chain sulfur atom of Cys208 in hydrogen bond formation is a distinctive structural feature that is presumed to confer substantially improved binding stability for the inhibitor-NDM-1 complex.

**Table 2.** Molecular interaction of Hydrogen bonds and Metallic coordinative bonds between NDM-1 and Antibiotics/Inhibitors.

| Ligand | Hydrogen bonds | Metal coordination bond |
| --- | --- | --- |
| Meropenem | His122 Asn220 | Zn1:His120 His122 His189 Ligand(1)<br>Zn2:Asp124 Cys208 His250 Ligand(2) |
| Hydrolyzed Meropenem | His122 Asp124 Lys211 Asn220(2) His250 | Zn1:His120 His122 His189<br>Zn2:Asp124(2) Cys208 His250 |
| Imipenem | Ala74 Asp124(2) Cys208 Lys211 Asn220(2)<br>His250 | Zn1:His120 His122 His189 Ligand(1)<br>Zn2:Asp124 Cys208 His250 |
| Hydrolyzed Imipenem | His122 Asp124 His189 Lys211 Asn220 His250 | Zn1:His120 His122 His189 Ligand(1)<br>Zn2:Asp124 Cys208 His250 Ligand(1) |
| RPX7009 | His122 His189 Asn220 | Zn1:His120 His122 His189 Ligand(1)<br>Zn2:Asp124 Cys208 His250 |
| QPX7728 | His189 Cys208 Lys211 Asn220 His250 | Zn1:His120 His122 His189 Ligand(1)<br>Zn2:Asp124 Cys208 His250 Ligand(1) |
| VNRX-5133 | Asp124 His189 Cys208 Lys211 Asn220 His250 | Zn1:His120 His122 His189 Ligand(1)<br>Zn2:Asp124 Cys208 His250 Ligand(1) |
| VNRX-5236 | Asp124 Lys211(2) Asn220 | Zn1:His120 His122 His189<br>Zn2:Asp124 Cys208 His250 Ligand(1) |

Previous studies have confirmed that residues His189 and His250 are critical for antibiotic substrate binding and hydrolysis^[27–28]^, and residues including Asp124, Asn220 and His250 are located in the vicinity of the NDM-1 active pocket (Fig.1). The catalytic center is vital for β-lactam hydrolysis, and interfering with these site interactions may block the enzyme’s antibiotic-degrading ability. Future inhibitor design targeting these key residues is expected to improve binding affinity and specificity, laying a solid foundation for subsequent studies.

**Fig 6.**
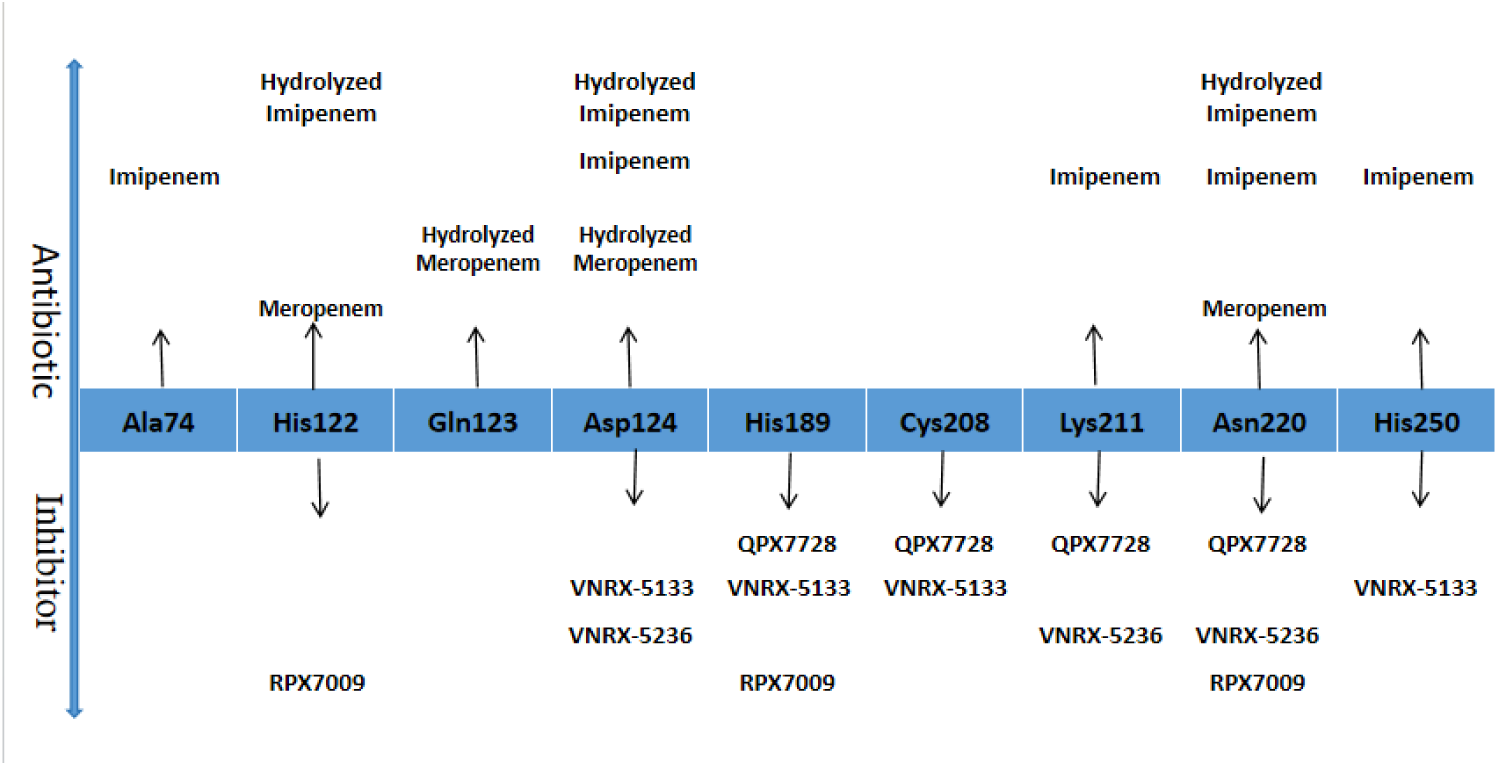
Hydrogen bond interaction between NDM-1 and antibiotics/inhibitors

### 3.3 FEL: Turning High-Dimensional Simulations into an Intuitive Energy Map

As a standard low-dimensional visualization paradigm for molecular simulation, the Free Energy Landscape (FEL) maps high-dimensional trajectory data onto collective variables, enabling unambiguous identification of stable conformations, transition states, and free energy barriers for quantitative energetic characterization of conformational distributions. The color gradient in the free energy landscape (FEL) diagram directly maps the corresponding free energy levels, with red denoting high-energy states and blue representing low-energy states.

In Figure 7-1, every enzyme-antibiotic binding complex has a low-energy region with a relatively concentrated spatial distribution, indicating a stable conformational state. Both meropenem and its hydrolysate occupy one distinct low-energy minimum, with the hydrolysate’s elongated well indicating limited conformational flexibility. The intact imipenem substrate has two interconnected low-energy regions, indicating it switches between multiple metastable states during the simulation, providing preliminary evidence for the inhibitor-induced fit mechanism. In contrast, hydrolyzed imipenem is mainly distributed in a broad single low-energy basin, with a relatively concentrated conformational distribution.

**Fig. 7-1.**
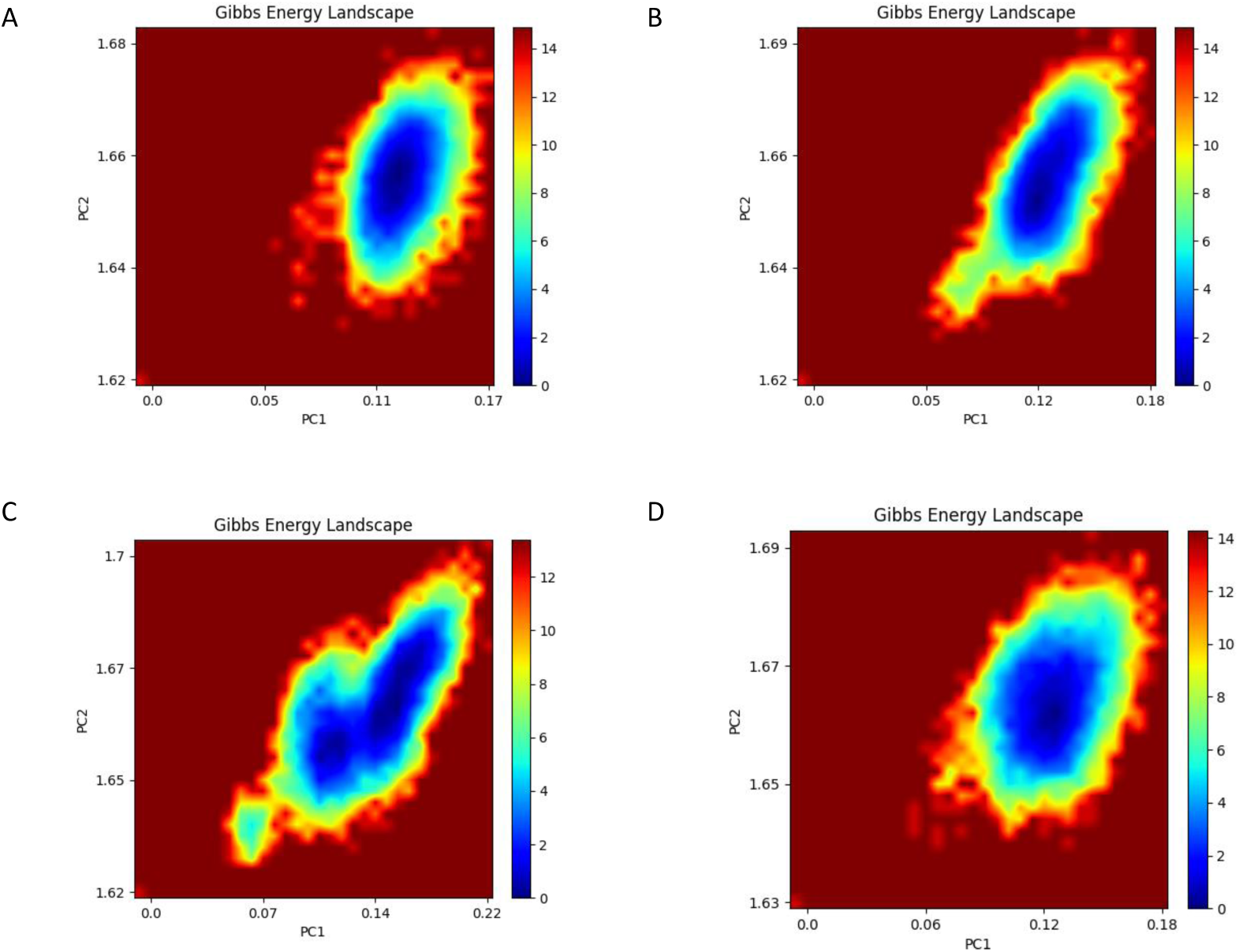
1. Free energy profile of NDM-1 and ligands (A) Meropenem; (B) Hydrolyzed meropenem (C) Imipenem; (D) Hydrolyzed imipenem Fig. 7 2. Free energy profile of NDM-1 with inhibitors (A) VNRX-5133; (B) QPX7728; (C) VNRX-5236; (D) RPX7009

**Fig. 7-2.**
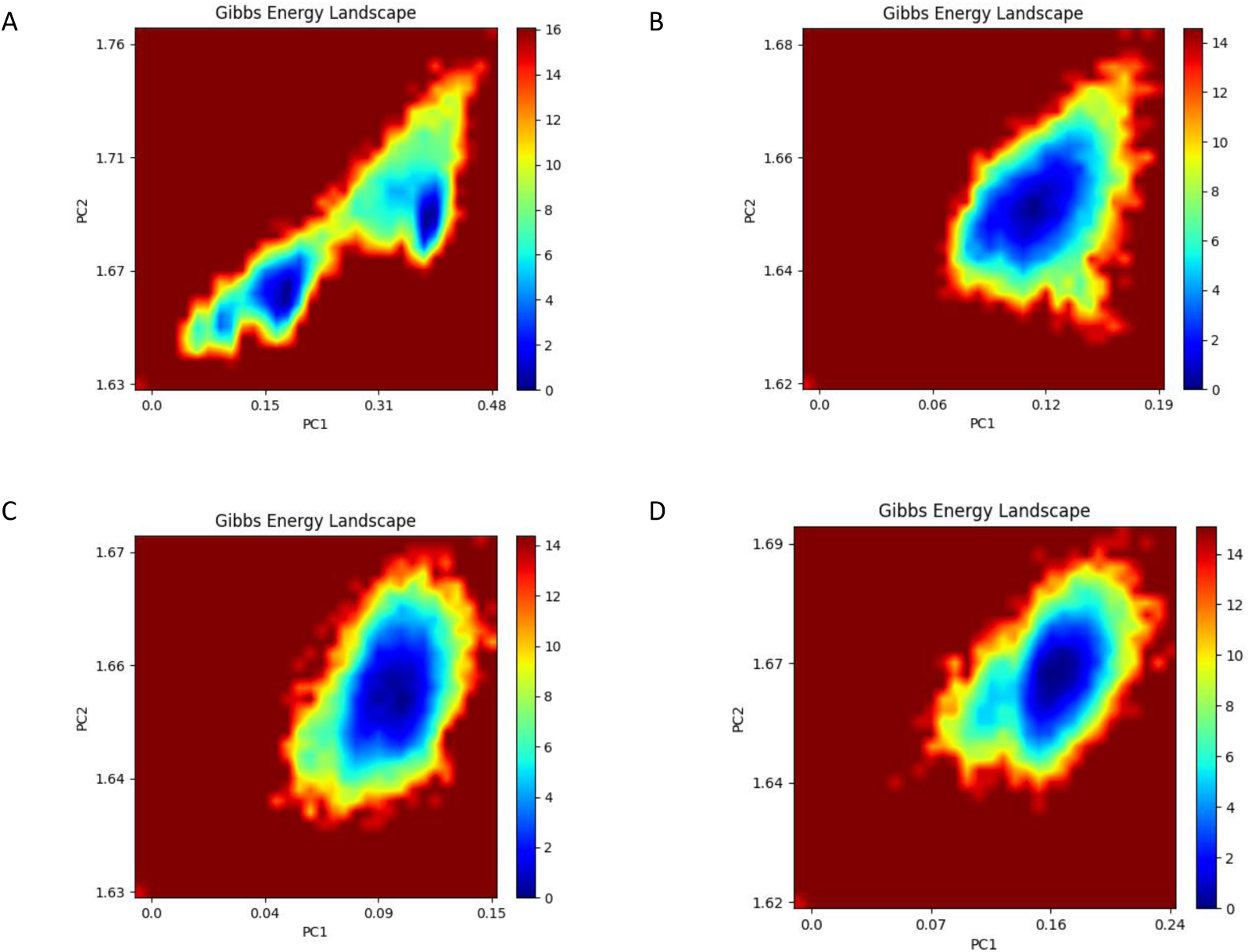
Free energy profile of NDM-1 with inhibitors (A) VNRX-5133; (B) QPX7728; (C) VNRX-5236; (D) RPX7009

Fig. 7-2 shows the free-energy landscape for the enzyme-inhibitor complex. Concentrated low-energy landscapes for QPX7728 and VNRX-5236 confirm persistent, stable dominant conformations across simulations. RPX7009 also shows a dominant low-energy region, with a slightly broader distribution range than the two materials above, yielding moderate stability. VNRX-5133 shows a dispersed low-energy region along the principal component axis, with two distinct low-energy states. The complexes fluctuate around one stable conformation during simulation, matching the rapid RMSD equilibration of their carbon skeletons, and confirming stable, well-defined binding modes. Notably, VNRX-5133 exhibits two low-energy basins corresponding to two energetically similar stable conformational states. This conformational diversity and adaptability likely arise from the boron inhibitor’s ability to interconvert between different hybridization states, improving its compatibility with various enzymes and providing preliminary evidence for the inhibitor’s induced-fit mechanism.

### 3.4 Bicyclic Boronates as Potent Transition State Analog Mimics

Bicyclic boronates are proven excellent transition state analog mimetics for SBLs and MBLs catalysis, due to their sp²-sp³ hybridization switchability and remarkable conformational restriction^[29–30]^. Molecular dynamics simulations reveal that cyclic boronic acid inhibitors markedly reduce the flexibility of the loop region adjacent to the NDM-1 active site, offering an important strategy for novel inhibitor design. VNRX-5133 and QPX7728, two representative bicyclic boronate inhibitors, have entered active clinical development in separate β-lactam combination regimens with cefepime and QPX2014^[31]^. Introducing additional conformational constraints at the carbon atoms adjacent to the boron atom rigidifies the bicyclic boronate moiety, potentially enabling the ligand to adopt a more optimal geometric configuration upon complexation with NDM-1.

## 4. Conclusion

In the present work, molecular docking and molecular dynamics simulations were employed to systematically characterize the intermolecular interactions between four cyclic boronic acid-based inhibitors (vaborbactam (RPX7009), taniborbactam (VNRX-5133), xeruborbactam (QPX7728), and ledaborbactam (VNRX-5236) and New Delhi metallo-β-lactamase 1 (NDM-1), with the hydrolysis of two representative carbapenem antibiotics (Meropenem and Imipenem) by NDM-1 set as the reference system. This study aims to combat drug resistance mediated by metallo-carbapenemases and restore the antimicrobial activity of carbapenem agents. The results show that these inhibitors potently inhibit NDM-1-catalyzed antibiotic hydrolysis via hydrogen-bond formation with critical amino acid residues in the enzyme active site. Notably, flexibility shifts in Loop3 and Loop10 of NDM-1’s secondary structure are part of the inhibitor-induced fit process. These conformational flexibility changes in Loop3 and Loop10 are part of the inhibitor-induced fit mechanism. Bicyclic boronic acid inhibitors display markedly superior binding affinity over monocyclic counterparts, with QPX7728 and VNRX-5133 being the most promising candidates identified to date. This study lays a solid theoretical foundation and provides actionable insights for the subsequent rational design of novel inhibitors based on the structure-function relationship of target proteins.

## Notes

### Competing Interest Statement

The authors have declared no competing interest.

